# Messages From the Small Intestine Carried by Extracellular Vesicles in Prediabetes: a Proteomic Portrait

**DOI:** 10.1101/2021.02.19.431856

**Authors:** Inês Ferreira, Rita Machado de Oliveira, Ana Sofia Carvalho, Akiko Teshima, Hans Christian Beck, Rune Matthiesen, Bruno Costa-Silva, Maria Paula Macedo

## Abstract

Extracellular vesicles (EVs) mediate communication in physiological and pathological conditions. In the pathogenesis of type 2 diabetes, inter-organ communication plays an important role in its progress and metabolic surgery leads to its remission. Moreover, gut dysbiosis is emerging as a diabetogenic factor. However, it remains unclear how gut senses metabolic alterations and whether this is transmitted to other tissues via EVs. Using a diet induced-prediabetic mouse model, we observed that protein packaging in gut derived EVs (GDE), specifically small intestine, is increased in prediabetes. Proteins related to lipid metabolism and to oxidative stress management were more abundant in prediabetic GDE compared to healthy controls. On the other hand, proteins related to glycolytic activity, as well as those responsible for the degradation of polyubiquitinated composites, were depleted in prediabetic GDE. Together, our findings show that protein packaging in GDE is markedly modified during prediabetes pathogenesis. Thus, suggesting that prediabetic alterations in small intestine are translated into modified GDE proteome, which are dispersed into the circulation where they can interact with and influence the metabolic status of other tissues. This study highlights the importance of the small intestine as a tissue that propagates prediabetic metabolic dysfunction throughout the body and the importance of GDE as the messenger.

## Introduction

Type 2 diabetes (T2D) is the most common chronic metabolic disease characterized by high blood glucose and it accounts for 95% of diabetes cases in adulthood [1]. The escalating incidence of T2D as a consequence of sedentary lifestyle and over-nutrition, imposes a substantial burden to health care systems worldwide. Among the therapeutic interventions, metabolic surgery which targets the small intestine, is the most effective procedure in diabetes remission [2;3]. Prediabetes is a high-risk state for T2D, and it is defined by glycemic levels higher than normal, but lower than diabetic limits [4]. This stage can be seen as a fork in the road, if left untreated, the likelihood for progression to diabetes is high. Importantly, with timely diagnosis and with patient’s compliance to lifestyle intervention, prediabetes can be reversed [5]. Obesity is a potent risk factor for diabetes. Moreover, a diet rich in fat and sugar, with large intake of finely processed grains and starchy carbohydrates is directly associated with this disease [6].

Blood glucose homeostasis requires a constant communication between insulin-secreting and insulin-sensitive organs. Failure in the coordination between these organs can lead to a rise in blood glucose levels and to prediabetes. Recently, extracellular vesicles (EVs) emerged as an important component of organismal communication and have been implicated in the pathophysiology of diabetes [7]. These small vesicles are able to transfer proteins and nucleic acids from a releasing cell to a distant target cell, thus modulating gene and protein expression and leading to functional changes [8–9]. Interestingly, visceral adipose tissue derived EVs trigger systemic inflammation, glucose intolerance and insulin resistance; hallmarks of prediabetes [10]. In obese patients, adipose tissue-EVs were able to modulate insulin responses in hepatocytes and muscle cells [11].

The intestinal system is daily exposed to food, millions of pathogens, and high concentrations of foreign antigens. In the case of the gut EVs, the composition and effects of EVs released by the resident microbiota have been explored [12–13]. For instance, these EVs are released into the gut mucosa and participate in local innate responses to invading bacteria through microbicidal activity [14] and mediate gut permeability through the regulation of tight junctions [15]. While the contribution of microbiota is crucial to maintain intestinal homeostasis, the epithelial cells are a critical component of the intestinal defense system forming a selective permeable barrier, allowing the absorption of nutrients, electrolytes and water, and blocking ingress of toxins and antigens. Epithelial cells in the gut were described to secrete EVs and up today they were described to play a role as antigen-carrying structures [16]. Whether EVs derived from the small intestine play a primordial role as a diabetogenic trigger needs to be elucidated. In fact, the relevance of the gastrointestinal tract in diabetes is elucidated by the fact that metabolic surgery improves glucose homeostasis more effectively than any known pharmaceutical or behavioral approach, causing durable remission in many patients with T2D [17]. Attempts to elucidate the exact mechanism by which gastro-intestinal surgery reverses T2D have implicated changes in the gut hormones, bile acid metabolism, intestinal nutrient sensing and metabolism, gut microbiota, and other factors [18]. Gut dysbiosis recently emerged as a possible player in the development of T2D [19]. Despite the literature on the impact of microbiota in T2D onset and development, the role of the small intestine EVs, independent of EVs derived from the biota is still elusive. While gut derived EVs (GDE) are considered to play an important role in intestinal diseases [20] it is not known how GDE protein content is modulated by a western diet. Therefore, it is timely to understand the importance of the small intestine as a metabolically active organ in prediabetes and the means by which it communicates with other tissues via EVs. Here we aim at characterizing the differences in the GDE proteome of diet-induced prediabetic mice compared to that of healthy mice fed standard chow.

## Results

### Description of the prediabetes mouse model

First, we established a mouse model that recapitulates many features of prediabetes [21]. High fat and sugar feeding leads to obesity, hyperinsulinemia and glucose intolerance. Therefore, we used an hypercaloric diet (HD) which composition is 58% fat and 18 % sucrose, in contrast with normal-chow diet (NCD) that is 11% fat with no added sugar. After 12 weeks of diet, HD mice demonstrate significantly increased body weight (Figure 1A, 1B) and an abnormal intrahepatic fat accumulation – a hallmark of liver steatosis (Figure 1C). They also developed glucose intolerance, evaluated by a glucose tolerance test in the previous week of sacrifice and correspondent area under the curve for each group (Figure 1D, 1E) (**Figure 1**).

**Figure 1.**
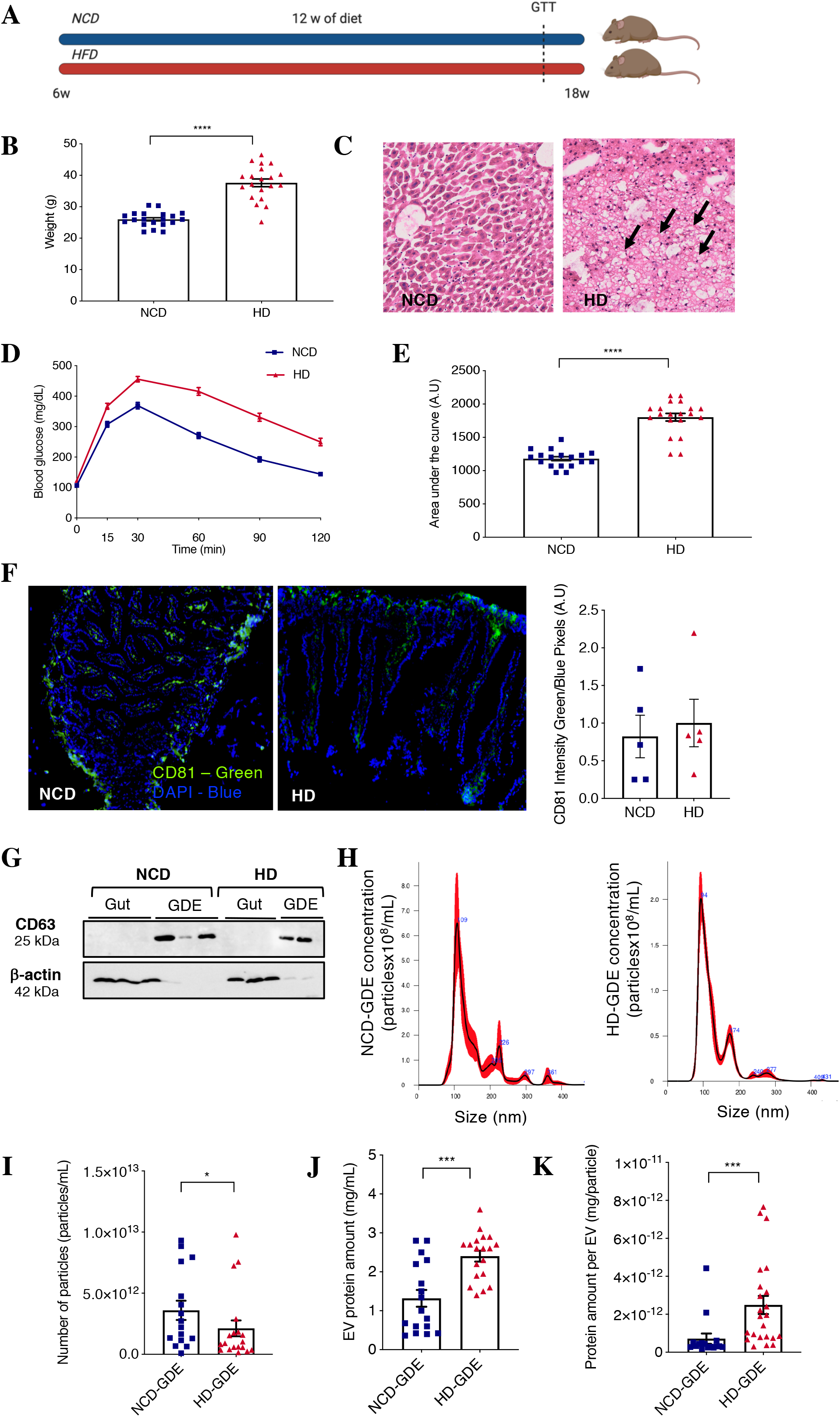
High fat diet induces prediabetes in mice and increases protein cargo in gut derived EV. **A**. Diet plan schematic representation. **B**. Body weight (n=20 per group). **C**. Representative Hematoxylin-Eosin images of liver histological sections of NCD and HD mice. White areas indicated by black arrows are lipid droplets, revealing steatosis in the liver of mice subjected to HD. **D.** Plasma glucose profile (md/dl) along a glucose tolerance test (GTT) at different time points after glucose bolus (0, 15, 30, 60, 90, 120 minutes) at week 17 of age. **E**. Mean area under the curve of the GTT. Results are expressed as mean ± SEM; n=20 per group. Unpaired t test with Welch’s correction for B and D; ****p<0.0001. **F**. Gut cross sections, EVs are labelled in green (CD81) and nuclei in blue (DAPI). NCD gut tissue in the left and HD gut tissue in the right. CD81 quantification in the gut. **G**. Immunoblotting of gut tissue and gut derived EV (GDE) isolated from NCD and HD mice. Protein extracts prepared from gut (10 μg) and from GDE (15 μg) were analyzed by western blotting using antibodies against EV marker CD63 and β-actin. **H**. Size and concentration distribution of GDE determined by nanoparticle tracking analysis (NTA). A representative NTA graph of GDE of mice fed with NCD (NCD-EV) (left) and HD (HD-EV) (right). **I**. Analysis of the number of particles per mL of sample. **J.** Protein quantification in GDE in mg per mL of sample, obtained by BCA. **K.** Protein content per GDE, represented in mg of protein per particle. Results are expressed as mean ± SEM; n=17 for NCD-EV and n=25 for HD-EV. Unpaired t test with Welch’s correction (F, I, J, K); ***p<0.0005, **p<0.005.

### Characterization of intestine derived extracellular vesicles

Once the prediabetic phenotype was validated in the diet-induced obese animals, we evaluated the presence of EVs in small intestine. We stained cross sections for CD81, a commonly used EV-marker, and the presence of this protein was detected in similar levels in the lumen of the intestine in both conditions (NCD and HD fed mice - Figure 1F). Moreover, western blot was performed to confirm the enrichment of CD63, a commonly used EV marker, in GDE from NCD and HD when compared with the tissue (Figure 1G). Next, GDE were characterized by nanoparticle tracking analysis and total amount of protein was quantified by BCA. The mean size of the particles, in both conditions, was around 100 nm (NCD=152.8±67.5; HD=128.6±50.7) (Figure 1H). While the number of EVs is decreased in HD animals (Figure 1I), protein cargo was higher in GDE isolated from HD-fed mice (HD-GDE) than in NCD-fed (NCD-GDE) mice (Figure 1J). Interestingly, the amount of protein per GDE was higher in prediabetic mice than in controls (Figure 1K) (**Figure 1**).

### Proteomics Study

EVs protein cargo provides valuable information regarding the cellular environment where they are produced [22]. We proposed that a diet induced dysmetabolic environment modulates the protein composition of GDE. Thus, to identify the impact of the diet-induced prediabetic gut milieu on GDE protein cargo, we isolated GDE (Supplementary Figure1) from mice under HD and respective controls and analysed it by mass-spectrometry (MS) (Supplementary Figure 2A). Potential contribution of gut bacteria EVs was ruled out by mass spectrometry analysis, which showed absence of bacterial proteins (Supplementary Figure 2B), and culture of gut conditioned media in antibiotics-free agar, which did not display bacterial growth (Supplementary Figure 2C).

### Bioinformatics analysis of the MS identified proteins

After filtering the identified proteins for reverse proteins used for false discovery rate and possible contaminating proteins (e.g. keratin and bovine proteins from cell culture media), we identified a total of 2467 proteins after removing isoforms from the same gene. Intensity based absolute quantitation (iBAQ) was calculated [23]. Excluding proteins with missing quantification values resulted in 2156 proteins (Supplementary file). The raw data was then normalized (Figure 2A). Subsequently, we distinguished between identified and regulated proteins. Regulated proteins are those at least two-fold regulated and with an adj. p-value < 0.05. After applying the aforementioned criteria to our data set, we obtained a list of robustly regulated 558 proteins, which shows that diet-induced prediabetes alters the protein composition of GDEs (Figure 2A). All data, before applying the filtering criteria, is represented in a volcano plot that describes the level of significance and magnitude of changes observed, comparing the HD with the NCD group (Figure 2B). The dotted line in the plot represents the threshold of adj.p-value<0.05, and from that line above each individual dot represents a protein within the 558 regulated ones.

**Figure 2.**
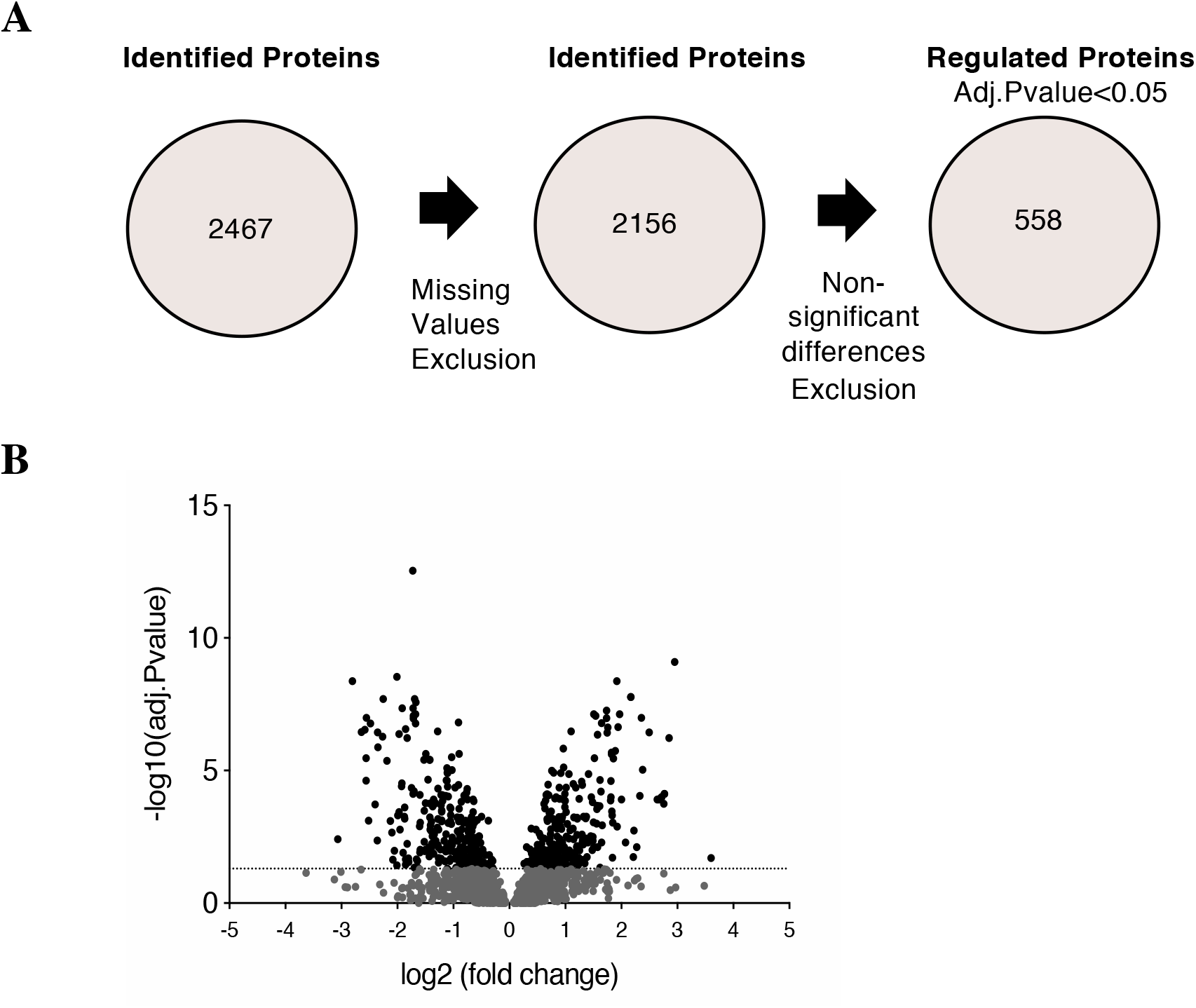
Study flow of proteomic approach for characterization of gut derived EV protein cargos. **A**. Strategy used for selection of proteins to be analyzed with confidence. **B**. Volcano plot representing the fold change and adj.p-value in the gut-derived EV (GDE) isolated from prediabetic mice and control groups. x-axis represents fold change, and y-axis adj.p-value. Dotted line sets the threshold of adj.p-value<0.05. In grey are the non-regulated proteins, in black the 558 regulated proteins.

### Gene Ontology

To gain insight into the biological role of the 588 regulated proteins, we performed Gene Ontology and Kyoto Encyclopedia for Genes and Genomes (KEGG) enrichment analysis, which revealed that the vast majority of proteins were related with metabolism. In terms of molecular function, enzyme activity (including isomerases, hydrolases and oxidoreductase) shows up in the top of list (Figure 3A). Most of the enriched pathways for gene ontology biological process emphasized lipid metabolism, highlighting fatty acids (FA) β-oxidation. Curiously, glycolysis-related and proteolysis in cellular protein catabolic processes-related proteins are mostly down regulated (Figure 3B). Considering KEGG pathways enrichment analysis, the affected proteins belong primarily to FA, carbohydrate and amino acid metabolism. Interestingly, alcoholism-related proteins show up with great significance (Figure 3C). In terms of down regulated proteins, based on the KEGG analysis the proteasome and focal adhesions proteins are markedly down regulated. Finally, to assess sub-cellular localization of the GDE protein cargo regulated by diet we used gene ontology cellular component analysis and found that the majority of altered proteins are mitochondrial, followed by the cytosolic ones. Once again, proteasome related proteins were down regulated (Figure 3D).

**Figure 3.**
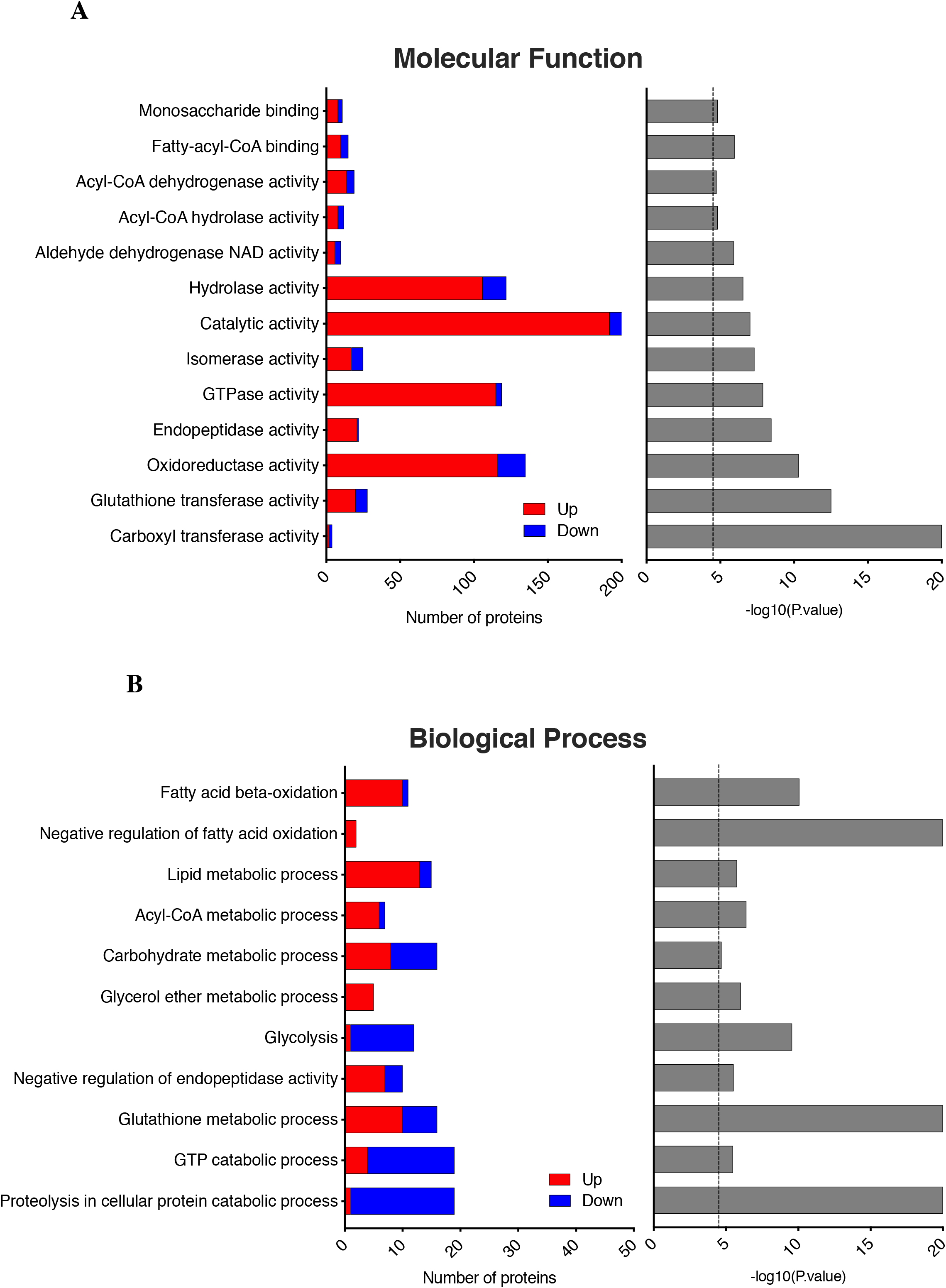
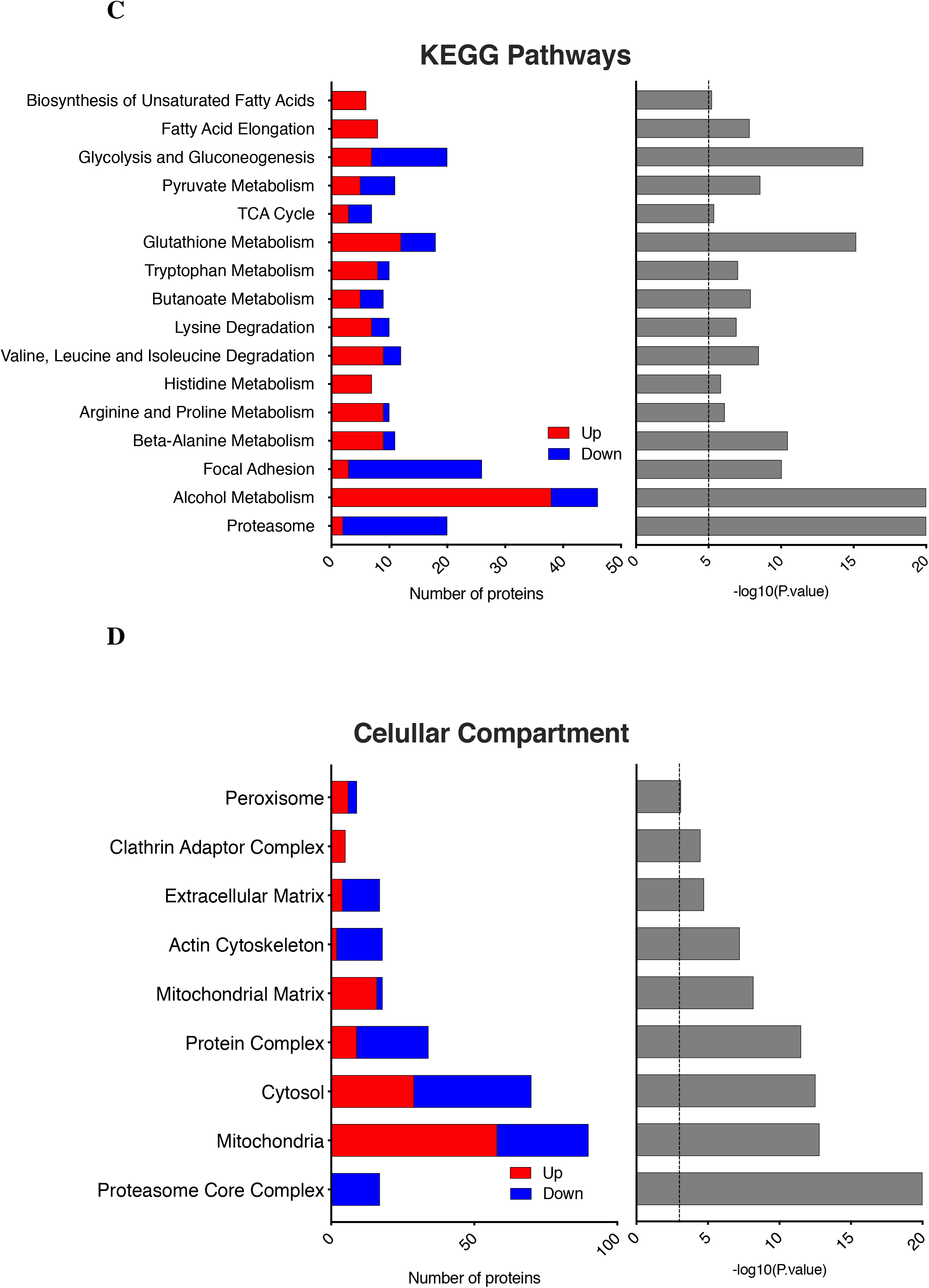
Characterization of prediabetic gut derived EV by GO and KEGG enrichment analysis directly against mouse annotation, according to GO terms. **A**. Molecular function analysis of the GDE identified proteins. GO enrichment analysis according to Molecular Function. **B**. Analysis of the main biological regulated by the proteins identified in the GDE. GO enrichment analysis according to Biological Process. **C**. KEGG enrichment analysis for KEGG pathway. **D**. GO enrichment analysis according to Cellular Compartment. On the left panels, changes are displayed as the number of proteins with increased (red) or decreased (blue) levels (horizontal axis). On the right panels, the horizontal axis indicates the significance -log10 (p-value) of the functional association, which is dependent on the number of submitted proteins in the class, the number of proteins annotated in the class, the total number of submitted proteins and the total number of mouse proteins.

### Cluster analysis for GDE regulated proteins

For a more detailed characterization of GDE proteome, we looked into the subset of proteins that were overly altered among groups. For that, we established an adj.p-value<0.0001 as cut-off (Figure 4A). The proteins with that significance are represented in the heatmap (Figure 4B). To gain insight into protein-protein interaction of GDE identified protein cargo, we used the STRING software to link some of the altered proteins and to display the level of relationship between them. The proteins that demonstrate higher level of interaction are the ones related with the proteasome, including both α subunits (PSMA) and β (PSMB). Actin related proteins, which function is to maintain cellular integrity, also display relevant interaction. Moreover, network analysis highlights strong tendency towards metabolic-related proteins (Supplementary Figure 2). The fold change of all altered proteins varied up to 2-fold when comparing HD with NCD GDE. Within metabolic pathways, the altered proteins are mainly involved in nutrient metabolism, such as glucose, FAs and amino acids. In addition, cell integrity, focal adhesion and proteasome-related proteins are all down regulated (Figure 4C and Supplementary Figure 4). Key GDE proteins up and down regulated by HD are also represented in their subcellular location (**Figure 5**). Noteworthy to mention, proteins related with carbohydrate metabolism, which ensures the supply of energy to the cells (PFKL, ALDOA, ALDOB, PGAM2, ENO1, PGM5), with the tricarboxylic acid (TCA) cycle (DLST, IDH1). cholesterol homeostasis (ACAT1, ACAT2) and proteasome (PSMA1, PSMA2, PSMA3, PSMA4, PSMA5, PSMA6, PSMA7, PSMB1, PSMB2, PSMB3, PSMB8, PSMB10) are down regulated by HD. On the other hand, proteins related with lipid homeostasis (ACOT1 ACOT2, ACOT3, ACOT5, ACOT6), FA mitochondrial β-oxidation (HDHA, ACADVL, ACADL, ECHS1, ACAA2), very-long chain FA peroxisomal β-oxidation (ECH1, ACOX1), amino acids metabolism (GSTA1, GSTA2) and antioxidant defence (ALDH2, ALDH1B1, PRDX6, GSTA4, GSTM2) are majorly upregulated. **Table 1** describes all the proteins up and down regulated according to their biological function.

**Figure 4.**
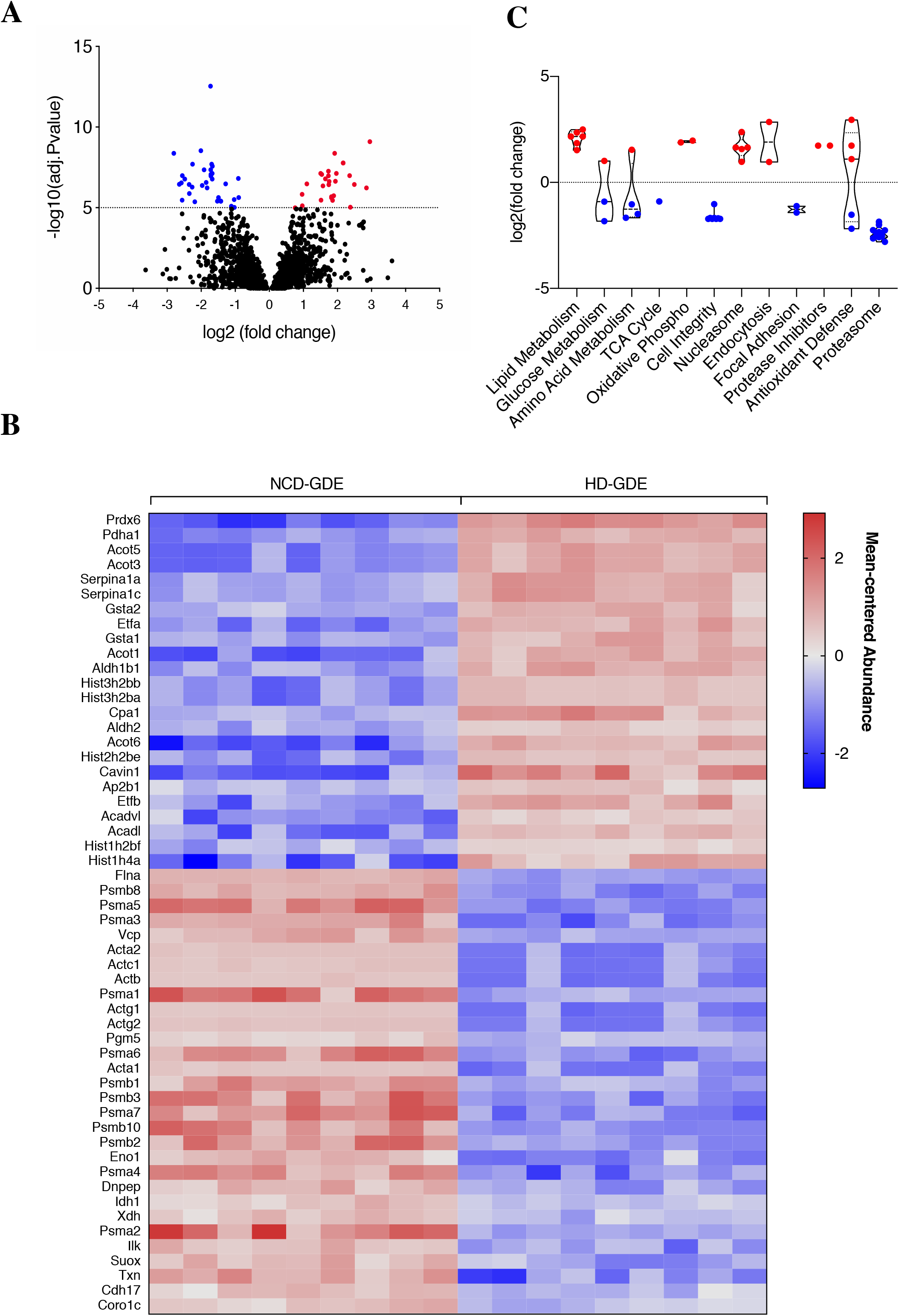
Cluster analysis of the top50 altered proteins of gut derived EV in prediabetic mice. **A**. Volcano plot representing the fold change and adj.p-value of gut-derived EV (GDE) proteins isolated from prediabetic mice and control groups. x-axis represents fold change, and y-axis adj.p-value. Dotted line sets the threshold of adj.p-value<0.0001, highlighting the most altered proteins in GDE from HD comparing with GDE from NCD. In blue are the most down regulated proteins and in red the most upregulated. **B**. Heatmap representing mean-centered abundance of the most altered proteins decorated in the previous volcano plot. **C**. Violin plot demonstrating the fold changes of the most altered proteins and correspondent cellular functions. Cellular metabolic process (GO) represented in green.

**Figure 5.**
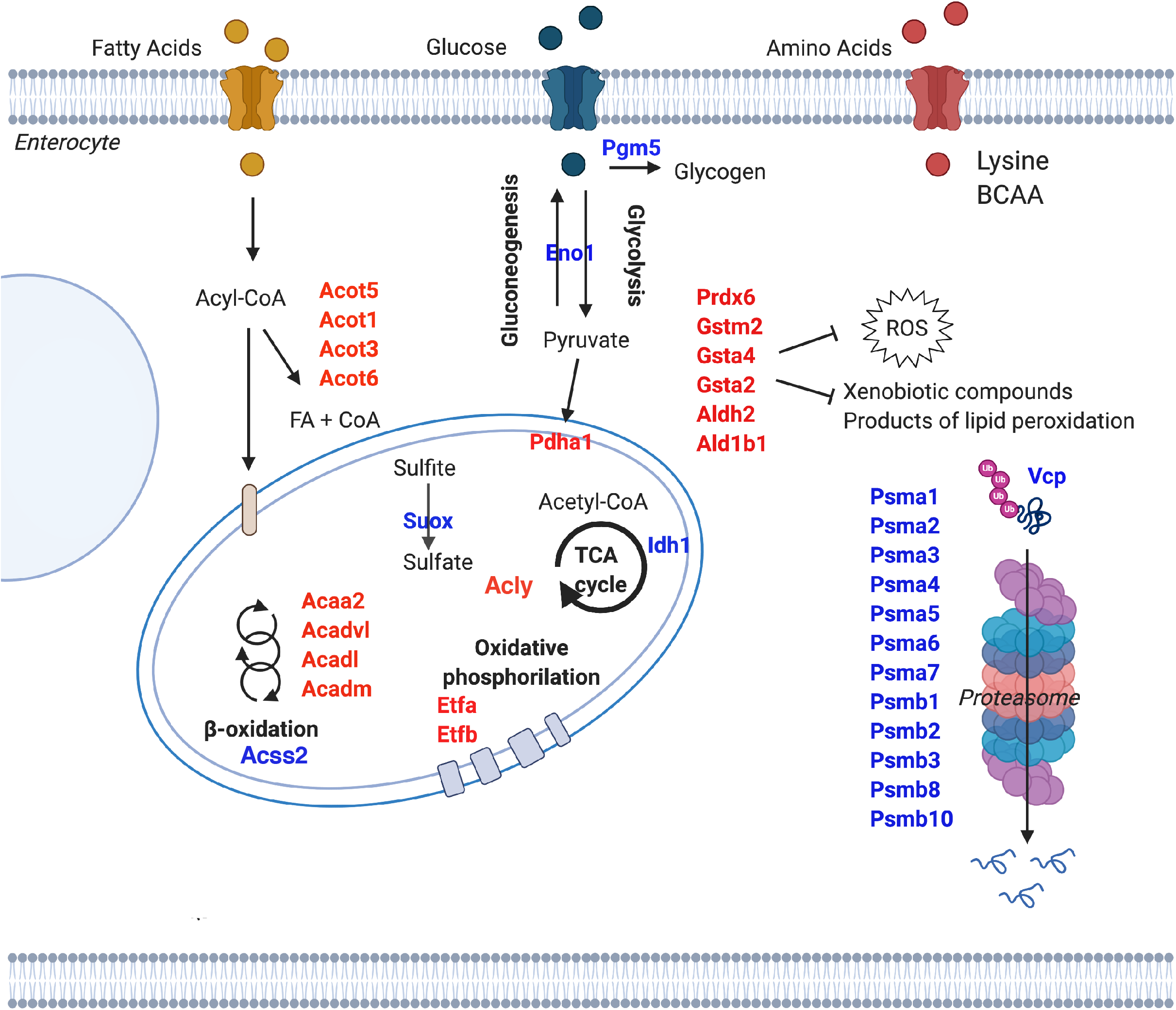
Overview of prediabetes impact on gut derived EV proteome. The identified proteins with higher differential expression are represented in the image according to their function and estimated subcellular localization. Red color for proteins upregulated and blue color for proteins down regulated in small intestine derived EV of mice fed with HD compared with controls, mice fed with NCD. ROS, reactive oxygen species; TCA, tricarboxylic acid cycle. Image done in BioRender.

**Table 1.** Biological function of the main up and down regulated proteins in HD-GDE compared with NCD-GDE by proteomics.

| Biological Function | Up | Down |
| --- | --- | --- |
| **Lipid Metabolism** |  |  |
| Mitochondrial β-oxidation | ACADVL;<br>ACADL | |
| Lipid Homeostasis | ACOT1;<br>ACOT2;<br>ACOT3;<br>ACOT5;<br>ACOT6 |  |
| **Glucose Metabolism** |  |  |
| Glycolysis |  | ENO1; PGM5 |
| Conversion of pyruvate to acetyl-CoA | PDHA1 |  |
| **Amino Acid Metabolism** |  |  |
| C-terminal amino acid release | CPA1 |  |
| Protein degradation |  | VCP |
| Peptide metabolism |  | DNPEP |
| Purine Metabolism |  | XDH |
| **TCA Cycle** |  | IDH1 |
| **Oxidative phosphorylation** | ETFA; ETFB |  |
| **Cell Integrity** |  | ACTA1;<br>ACTA2; ACTB;<br>ACTC1;<br>ACTG1;<br>ACTG2;<br>CORO1C;<br>FLNA |
| **Nucleosome** | HIST1H4A;<br>HIST3H2BB;<br>HIST3H2BA;<br>HIST2H2BE;<br>HIST1H2BF |  |
| **Endocytosis** |  |  |
| Caveolar endocytosis | CAVIN1 |  |
| Clathrin-dependent endocytosis | AP2B1 |  |
| **Focal adhesion** |  | ILK; CDH17 |
| **Protease inhibitor** | SERPINA1A;<br>SERPINA1C |  |
| **Antioxidant Defense** | PRDX6;<br>GSTA4;<br>GSTA1;<br>GSTA2;<br>ALDH2;<br>ALDH1B1 | TXN; SUOX |
| **Proteasome** |  | PSMA1;<br>PSMA2;<br>PSMA3;<br>PSMA4;<br>PSMA5;<br>PSMA6;<br>PSMA7;<br>PSMB1;<br>PSMB2;<br>PSMB3;<br>PSMB8;<br>PSMB10 |

## Discussion

The relationship between diet composition and the incidence of metabolic disorders is a public health concern. One of the first organs and by far the largest one to contact with food and its derivates is the gut, as a consequence it is where the first organismal response should be found [24]. Here, we report that GDE isolated from animals fed an HD, mimicking the western one, have a distinct proteome that strongly reflects the intestinal dysmetabolic milieu. We found numerous pathways associated with macronutrients altered in GDE isolated from HD-fed mice; of great relevance the ones related with lipidic and carbohydrate metabolic processes.

From the enrichment analysis we observed that both glycolysis-gluconeogenesis and pyruvate pathways are within the most affected ones (Figure 3B). Rate limiting enzymes for glycolysis were found to be less abundant, namely hexokinase (HK) and phosphofructokinase (PFKL), thus suggesting changes in glucose utilization and lactate production. In accordance, in diabetic rats, phosphorylation of glucose is inhibited in tissues such as the heart, skeletal muscle and liver [25; 26; 27; 28]. Interestingly, in hearts from diabetic rats, PFK and HK down regulation is reverted by an oral anti-diabetic drug - metformin [29]. We suggest that the observed decreased levels of PFKL in prediabetic GDEs is reflecting a similar down regulation in the small intestine of prediabetic mice and is an early event in the progression of the disease [30]. Down in the glycolysis pathway, pyruvate dehydrogenase A1 (PDHA1), which belongs to the PDH complex, was upregulated in HD GDEs, indicating a possible compensation in the pathway given the importance of acetyl-CoA for the TCA cycle and for many other proteins and lipidic biochemical reactions. PDH is fundamental for metabolic flexibility, the ability of switching from carbohydrates to lipid fuel as energy sources, and the inability of cells to do so is an important hallmark of T2D [31]. Our results suggest that in the early stages of the disease gut cells tackle the insult of HD feeding by changing the source of fuel. The decrease in glycolysis might be explained by the low amount of fibers provided by the diet; since the percentage of fibers in HD is considerably reduced and in part replaced by sucrose. Likewise, fibers have a beneficial effect on NCD mice, by improving intestinal integrity, enhancing energy expenditure and reducing inflammation [32]. In fact, because HD has increased sucrose, we envisage that sucrose is being directed to an alternative pathway, such as de novo lipogenesis, known to also occur in the gut [33].

In accordance with the high lipidic content present in the HD, we observed that lipid metabolism (biosynthesis of unsaturated FA and FA elongation) is vastly altered. The excess of fat in the diet is known to cause increased fatty acids uptake, which can be either metabolized for the production of energy or, when in excess, will be stored. Several proteins involved in mitochondrial β-oxidation (HADHA, ACADVL, ACADL, ECH1, ECHS1), which generates acetyl-CoA to feed the TCA cycle, and peroxisome β-oxidation (ECH1, ACOX1), major fatty acids degradative pathway, are upregulated in GDE by HD, suggesting that fatty acids are favoured as energy source over glucose (Figure 5). In addition, over-activation of lipid metabolism related pathways is a common feature of obesity and T2D, leading to a state of hyperglycemia [34; 35; 36]. Moreover, a family of acyl-CoA thioesterases (ACOTs) were particularly upregulated; these are intermediaries in the decision whether fatty acids are directed to the TCA cycle or stored, (Figure 5, Supplementary Figure 4) [37; 38]. Altogether, this suggests that HD shifts the burden of energy provision from carbohydrates toward fat derived from diet, since ACSS2 (a protein that conducts fatty acids to the TCA cycle) is down regulated and PDHA1 (which provides a link between glycolysis, TCA cycle and lipogenesis) is upregulated in order to ensure the energetic need of the cell. Based on recent literature, obesity-induced activation of ACOTs directs fatty acids towards triglycerides synthesis for incorporation into chylomicrons or VLDL particles [39; 40]. Noteworthy, some studies consider activation of ACOTs protective against fatty acids over-supply, slowing the flux of fatty acids to downstream metabolic pathways [41; 42; 43]. Whether this is the case in the gut, needs to be further explored. Additionally, intestinal cells in the HD model appear to prioritize production of energy from fatty acids, instead of carbohydrates. This is shown by the upregulation of ATP-citrate lyase (ACLY), an enzyme responsible for the conversion of citrate to acetyl-CoA – the first step in DNL. Overall, the major enzymes found altered within the gut EVs are very much responsible for regulating acetyl-CoA levels in the cells, given that this molecule is a fundamental node in metabolism, in the future it would be of major interest to quantify it in enterocytes.

Increased fatty acids oxidation promotes mitochondrial dysfunction, as well as fatty acids peroxidation and glucose dysmetabolism, altogether leading to reactive oxygen species (ROS) accumulation to pernicious levels [44]. Oxidative stress is implicated in various diseases including diabetes [45]. We observed several proteins involved in antioxidant defense mechanisms, such as removal of free radicals (PRDX6) and elimination of xenobiotic compounds and products of lipids peroxidation (GSTA4, GSTM2) upregulated in HD-GDE, suggesting an activation of an antioxidant response in the gut cells (Figure 5). Consistent with our model, the presence of numerous isoforms of the aldehyde dehydrogenase (ALDH) family was upregulated in HD-derived GDE. ALDHs are a family of oxidizing enzymes responsible for cellular detoxification by oxidation of biogenic and xenogeneic aldehydes that accumulate through metabolism [46]. In particular, we highlight ALDH1B1 due to its ability to metabolize acetaldehyde and maintain glucose homeostasis [46]. Of interest, ALDH2 has been explored as a pharmacological target, due to its role in cellular damage defense against oxidative stress induced by pathological conditions, for instances high glucose [47].

Besides glucose and fatty acids, amino acids, in particular lysine, also play important roles in providing energy to gut epithelial cells (Figure 3C) [48]. Lysine, which pathway we see affected in HD-GDE, mediates protein biosynthesis, such as of carnitine, a key factor in fatty acid metabolism [49]. Interestingly, carnitine deficiency due to poor lysine diet induces triglycerides accumulation [50]. Evidence that the HD not only induces obesity, but also a general pathological state arises from the massive alteration of proteasome-related proteins. The proteasome is recognized as an important target in many diseases [51]. Our data showed down regulation of proteasome proteins, which is consistent with decreased proteasome function reported in a broad array of chronic diseases (Figure 3C; 5) and defective degradation of damaged proteins, a hallmark of neurodegenerative diseases [52]. One hypothesis is that the decreased content of proteasome proteins in prediabetic GDE mirror in the cell of origin. Alternatively, the GDE producing cells can be consuming proteasome to counteract the deleterious effects of oxidative stress, resulting in lower packaging of proteasome proteins in GDEs.

The present study is the first comprehensive analysis of GDE protein content. We found that diet composition strongly modulates GDE proteomic profile. Several lines of evidence support a model in which the metabolic alterations induced by the HD are translated in a similar manner in alterations in GDE protein cargo. These results bring light into the important modification taking place in the small intestine and how it translates into GDE content. We propose that future studies should focus on whether GDE can act in local and distant cellular targets and mediate prediabetes. We also propose that the analysis of GDE in biofluids may represent a strategy to non-invasively detect metabolic modifications in the gut linked with prediabetes and potentially be employed early diagnose and follow-up prediabetes patients.

## Material and methods

### Animals

Male C57Bl/6J mice were housed in a temperature-controlled room, in a 12-hour light/dark cycle. For induction of prediabetic phenotype, C57BL/6J mice started on a high-fat diet at 6 weeks of age, with free access to food and water, for 12 weeks. Hypercaloric diet (HD) (OpenSource Diet, D12331) composition is 16.5% protein, 25.5% carbohydrate, and 58% fat with 13% addition of sucrose. The control normal chow diet (NCD) (Special Diets Services, RM3) composition is 26.51% proteins, 62.14% carbohydrates and 11.35% fat and no sucrose. At 18 weeks of age, 12 weeks of diet, mice were sacrificed. Mice were anesthetized using 2% isoflurane. Mice were sacrificed under 12 hours of fasting, with a previous feeding period of 2 hours, with the previous 8 hours of fasting. Blood, liver and small intestine were collected from prediabetic mice and respective controls. All animals were treated according with National and European Union Directive for Protection of Vertebrates.

### Glucose Tolerance Test

At week 11 of diet GTT was performed. Mice were fasted overnight and weighed in the morning. Mice were then given an intra-peritoneal injection of a glucose solution (20% m/v – Sigma Aldrich) at 2g/kg body weight. Serum blood glucose levels were measured at 15, 30, 60, 90 and 120 min after injection. Blood glucose levels we measured using OneTouch Ultra glucose meter (LifeScan Inc).

### Small Intestine Histology

After sacrifice, liver and small intestine were washed in PBS and fixed for histology and a portion of small intestine was snap frozen in liquid nitrogen for later analysis by western blot. The clean gut was placed in 2% Paraformaldehyde with 20% Sucrose, overnight at 4°C; washed three times in Phosphate Buffered Saline (PBS), 10 min each; transferred to 30% sucrose for 2 hours at 4°C; left overnight at 4°C in Optimal Cutting Temperature compound (OCT) with 30% Sucrose (1:1). In the following day, tissues were embedded in 20% Sucrose (1:4) + (OCT) (3:4) in the presence of dry ice and immediately stored −80°C. Consequently, tissues were sliced into 6 μm thin sections in the cryostat and stored at −80°C for following applications. Gut samples were used for CD81 staining and liver samples for Hematoxylin-Eosin staining.

### Exosomes Isolation and Nanoparticle Tracking Analysis

After sacrifice the small intestine was removed, washed with PBS and cultured overnight, at 37°C and 5%CO2, in RPMI medium (Sigma) supplemented with 1% Penicillin/Streptomycin and 10% Fetal Bovine Serum EV-free, to avoid contaminations from serum EVs. In the following day medium was collected. Conditioned medium was submitted to two initial centrifugations (10 min, 500 g and 20 min, 3 000 g) to remove any suspended or dead cells in the medium. To remove large EVs, media was centrifuged (20 min, 12 000 g) and the pellet was discarded. The supernatant enriched in small EVs was again centrifuged (2 h 20 min, 100 000 g), and the EVs enriched pellet was collected. For sucrose cushion purification, this pellet was resuspended in 14 mL filtered phosphate buffered saline (PBS, Corning 15313581, NY, US) and added the top of 4 mL sucrose solution (D2O containing 1.2 g of protease-free sucrose and 96 mg of Tris base adjusted to pH 7.4). A new ultracentrifugation was performed (1 h 10 min, 100 000 g), after which 4 mL of the sucrose fraction was collected using a 18 G needle placed at the bottom of the ultracentrifugation tube (away from the pellet). Finally, 16 mL of PBS were added to the collected sucrose/EVs solution and an overnight (16 h, 100 000 g) ultracentrifugation was performed. The pellet containing the isolated EVs was resuspended in filtered PBS. The concentration and size of EVs were analyzed in NanoSight NS300 (NS3000) system equipped with a blue laser (405 nm), according to the manufacturer’s instructions. EVs were diluted 1:1000 in filtered sterile PBS. Each sample analysis was conducted for 90 seconds and measured 5 times using Nanosight automatic analysis settings. The software calculated the size distribution in nanometers and the concentration in number of particles per mL.

### Western Blotting

Protein extracts for western blot analysis were obtained using cold lysis buffer (20 mM Tris- HCl pH 7.4, 5 mM EDTA pH 8.0, 1% Triton-X 100, 2 mM Na3VO4, 100 mM NaF, 10 mM Na4P2O7) in the presence of protease inhibitors (completeTM, Mini, EDTA-free Protein inhibitor cocktail tablets, Roche, Sigma). Gut tissue and GDE were homogenized in the lysis buffer and sonicated three time for 30 seconds each at 10 μm amplitude, incubated on ice between each sonication step. Lysates were centrifuged at 18.000 g for 10 min at 4°C. Soluble fraction was collected and protein concentration was determined using the PierceTM BCA Protein Assay kit (Thermo Fisher). For protein expression analysis, equal amounts of total protein (20 μg) were mixed with sample buffer (250 mM Tris-HCl pH 6.8, 8% SDS, 40% glycerol, 8% β-mercaptoethanol, bromophenol), before being denaturated at 95°C for 10 min. Samples were then resolved by an 8% sodium dodecyl sulfate-polyacrylamide gel electrophoresis (SDS-PAGE). Proteins were subsequently electrotransferred to a polyvinylidene fluoride (PVDF) membrane (Immobilon-P Membrane, PVDF, Millipore) for 20 min at 25 mA and 2,5 mV in a Trans-Blot Turbo TM Transfer System (BioRad). Membranes were blocked for 1 hour at RT with 3% BSA (Thermo Fisher) in tris-buffered saline with 0,1% Tween (TBS-T) (blocking solution). Membranes were incubated overnight at 4°C, with CD63 antibody (Santa Cruz Biotechnology) prepared in 3% BSA in TBS-T, diluted 1:1000. Membranes were washed three times with TBS-T and incubated for 1 hour at RT with the secondary antibody anti-rabbit antibody diluted 1:5000 (Santa Cruz Biotechnology). Blots were developed with ECL (ECL Prime, GE Haelthcare) according to manufacturer’s instructions. Chemidoc Touch (Biorad) was used to detect chemilumunescence. Band intensities were quantified usinf ImageLab software and normalized by β-actin as a loading control.

### Nano-LC-MSMS analysis

Peptide samples were analysed by nano-LC-MSMS (Dionex RSLCnano 3000) coupled to a Q-Exactive Orbitrap mass spectrometer (Thermo Scientific). Briefly, 5 μL of sample was loaded onto a custom made fused capillary pre-column (2 cm length, 360 μm OD, 75 μm ID) with a flow of 5 μL per min for 7 min. Trapped peptides were separated on a custom made fused capillary column (20 cm length, 360 μm outer diameter, 75 μm inner diameter) packed with ReproSil Pur C18 3-μm resin (Dr. Maish, Ammerbuch- Entringen, Germany) with a flow of 300 nL per minute using a linear gradient from 92 % A (0.1% formic acid) to 28 % B (0,1% formic acid in 100 acetonitrile) over 93 min followed by a linear gradient from 28 % B to 35 % B over 20 min at a flowrate of 300 nl per minute. Mass spectra were acquired in positive ion mode applying automatic data-dependent switch between one Orbitrap survey MS scan in the mass range of 400 to 1200 m/z followed by HCD fragmentation and Orbitrap detection of the 15 most intense ions observed in the MS scan. AGC target value in the Orbitrap for MS and MSMS scans were 1,000,000 ions at a resolution of 70,000 at m/z 200 for MS scans, and 25,000 ions at m/z 200 for MSMS scans. Peptide fragmentation in the HCD cell was performed at normalized collision energy of 31 eV. Ion selection threshold was set to 5,000 ions for MSMS analysis, and maximum injection time was 100 ms for MS scans and 200 ms for MSMS scans. Selected sequenced ions were dynamically excluded for 60 seconds.

### MSMS analysis

Mass accuracy was set to 5 ppm on the peptide level and 10 Da on the fragment ions. A maximum of four missed cleavages was used. Carbamidomethyl was set as a fixed modification. M oxidation, N-terminal protein Acetyl, Q Deamidation and N Deamidation were set as variable modifications. The MSMS data was searched against all reviewed mouse proteins from UniProt with concatenation of all the sequences in reverse maintaining only lysine and arginine in place. The data was searched and quantified with both MaxQuant [53] and VEMS [54].

### Statistical Analysis

Data are presented as means ± SEM. GTT curves, bar plots, volcano plots, heatmap and linear regression analyses were done using GraphPad Prism 8 (GraphPad Software). Differences significance was calculated through unpaired student’s t tests with Welch’s correction; one-way ANOVA followed by Tukey-Kramer multiple comparison tests. Differences were accepted as statistically significant at p<0.05.

## Author’s contributions

IF and RMO carried out the animal model of disease, western blotting and drafted the manuscript. IF did the isolation of the extracellular vesicles and characterization by NTA; gut histology and performed bioinformatics analysis, together with MS, for graphical representation. ASC prepared GDE for MS analysis and RM was responsible for the mass spectrometry analysis and bioinformatics. Mass spectrometry was performed by HCB. MPM and BCS conceived the study, participated in its design and coordination and edited the manuscript. All authors read and approved the final manuscript.

## Acknowledgements

This work was supported by the Fundação para a Ciência e Tecnologia (Grant No. PTDC/MEC-MET/29314/2017) and Programa Gilead GÉNESE - Edição de 2019 awarded to MPM, RMO and BCS. IF is a recipient of Fundação para a Ciência e Tecnologia fellowship (PD/BD/114044/2015). AT is a recipient a European Foundation for the Study od Diabetes and Japonese Diabetes Society Reciprocal Travel Research Fellowship Programme.

## Supplementary material

**Supplementary Figure 1 Study flow of proteomic approach for characterization of gut derived EV protein cargos**

**A**. Schematic representation of the detailed protocol used for isolation of GDE. Image done in BioRender.

**B.** Schematic representation of the experimental process of analysis of gut derived EV (GDE). Image done in BioRender. **C**. Bar plot indicating the number of proteins uniquely identified from different species when searched against all proteins from all species in UniProt. **D**. Picture of LB-agar (without antibiotics) petri dish after 24 after hours of inoculation with media where guts were deposit overnight at 37°C prior to EVs isolation.

**Supplementary Figure 2 Protein-protein interaction network**

The network was built using the STRING online software with a medium confidence level (0.4). Each circle represents one protein. The line thickness indicates the strength of the evidence, with thicker connections indicating higher confidence in the protein-protein interaction. Green circles represent the proteins belonging to cellular metabolic process according to GO enrichment analysis for biological processes.

**Supplementary Figure 3 Individual protein alteration levels according to its biological function**

Data plots demonstrating the log fold changes of each protein in the image in accordance to its related pathway.

